# CDC42 Inhibitors Alter Patterns of Vessel Arborization in Skin and Tumors in vivo

**DOI:** 10.1101/2024.04.09.588609

**Authors:** Linh M Vuong, Stephanie Hachey, Jessica Shiu, Danny F Xie, Noel Salvador, Nicoletta Brindani, Sine Mandrup Bertozzi, Maria Summa, Rosalia Bertorelli, Andrea Armirotti, Rachel Pham, Vance SH Ku, Terry Nguyen, Bernard Choi, Christopher CW Hughes, Marco De Vivo, Anand K Ganesan

## Abstract

Tumors that arise in the epidermis must develop a vascular supply to grow beyond a millimeter in depth. This process requires CDC42 GTPases such as CDC42, RhoJ and RhoQ. Despite this dependence on angiogenesis for growth, melanoma tumors are minimally responsive to current anti-angiogenesis agents, highlighting the need for more effective drugs in this class. Here we integrate antibody infusion, optical tissue clearing, multiphoton imaging, and three-dimensional semi-automated tracing to develop a quantitative approach to measure changes in vascular architecture in skin and skin tumors. This new approach uncovered differences in vessel arborization in the skin of RhoJ KO mice as compared to wild-type mice. Furthermore, novel small molecules that inhibit CDC42 GTPases inhibited both tumor growth and vessel branching within tumors to a similar degree as Braf inhibitors, which are commonly used to treat melanoma. In contrast to Braf inhibitors, however, which only affected tumor vasculature, CDC42 inhibitors affected vascularization in both tumor and normal skin without apparent toxicity to endothelial or stromal cells. These novel CDC42 inhibitors similarly blocked vessel branching in human cell-based micro-physiological models of normal and tumor vessels. RNA sequencing revealed reduced expression of multiple angiogenesis-related genes in drug-treated skin. Taken together, these studies identify a new class of pharmacologic agents that inhibit vessel branching in both normal skin and tumors with potential utility for treating skin cancer and skin diseases characterized by pathologic angiogenesis.

## Introduction

Vascular morphogenesis in tissues is a dynamic process where endothelial cells modulate their cytoskeleton and cell matrix adhesions to properly respond to environmental cues (*1*). The dynamic rearrangement of endothelial cells’ cytoskeleton allows them to migrate both forward and backward within the growing blood vessel (*2–4*). Generation of the capillary plexus depends on the regression of existing vessels during the maturation process, and similar types of vessel regression are observed in conditions where the adult vasculature needs to be remodeled (*5*). All tumors, particularly those that arise in the epidermis that lack their own vasculature, are dependent on the ability to induce the growth of vessels around them (*6–8*). Tumors rapidly accelerate angiogenesis by producing a cornucopia of angiogenesis-promoting factors, inducing a haphazard, tortuous vessel tree that is unable to deliver nutrients evenly to the tumor (*9, 10*).

Combining tissue clearing with semi-automated tracing has provided great insight into how angiogenesis occurs in the skin and other tissues (*11–13*). This approach has been recently applied to tumors (*14, 15*). The tumor vasculature is characterized by large tortuous vessels with leaky basement membranes and unperfused voids (*16, 17*). While longitudinal imaging approaches have provided structural insight into how vessels develop or repair after injury (*5, 18*), this information alone has not identified better therapies for conditions characterized by pathologic angiogenesis, such as inherited skin vascular anomalies (*19, 20*) or melanoma tumors (*6*), where existing anti-angiogenic therapies are many times toxic or only minimally effective (*21*).

Many intracellular signaling pathways are known to activate angiogenesis (*22*). Vascular endothelial growth factors, which activate intracellular signaling cascades that induce actin remodeling and focal adhesion assembly (*23*), are major chemo-attractants for migrating endothelial cells. For example, spatial activation of VEGFR2 signaling is critical in the maintenance of the adult vasculature (*5*). Semaphorins have also been identified as regulators of vasculogenesis – Semaphorin 3E binds to the PlexinD1 receptor to induce receptor internalization, focal adhesion disassembly, and, subsequently, vessel regression (*24, 25*). Two CDC42 GTPases, RhoJ and CDC42, are critical mediators of both attractive and repulsive cues generated by activation of VEGF and Plexin receptors, respectively (*26*). RhoJ, a protein with 55% homology to CDC42, is highly expressed in endothelial cells and induces actin depolymerization, focal adhesion disassembly, and endothelial cell contraction (*27, 28*). CDC42, in contrast, activates focal adhesion assembly and stimulates the migration of endothelial cells (*29–32*). RhoJ and CDC42 mediate crosstalk between receptors that activate angiogenesis, particularly the VEGFR2 and Plexin receptors (*28*). RhoJ deficiency results in impaired VEGF-induced endothelial cell migration (*33*), impaired retinal vascular angiogenesis early during development (*34, 35*), and impaired tumor angiogenesis (*36*). Targeting RhoJ/CDC42 with small molecules has been problematic because of their globular structure and lack of readily apparent druggable pockets (*37, 38*). We recently used a structure-based drug design approach to discover a class of RhoJ/CDC42 inhibitors that block downstream effector interactions (*39, 40*). These agents blocked S6 and ERK activation, inhibited angiogenesis in vascularized tumor organoids, and inhibited the growth of mouse and patient-derived tumors *in vivo* (*39, 40*).

Here we sought to determine how RhoJ/CDC42 and drugs that inhibit their signaling modulate angiogenesis *in vivo*. We developed a platform combining tissue clearing and vascular labeling with fluorescent antibodies and three-dimensional semi-automated vessel tracing and discovered that RhoJ deficient mice had significantly altered vessel arborization in their skin with no other signs of overt toxicity. We applied this new imaging approach to image patient-derived melanoma xenografts treated with a recently discovered family of CDC42 inhibitors and other standard-of-care melanoma therapies (BRAF inhibitors), demonstrating that both agents had similar tumor suppressive effects. Both agents also inhibited blood vessel branching and the accumulation of vessel endpoints. These findings were confirmed *in vitro* using humanized organotypic micro- physiological systems (VMO and VMT) that recapitulate *in vivo* neo-vascularization in normal tissues and tumors. In contrast to vemurafenib, which induces necrosis of tumor cells with subsequent effects on vessels, we discovered that CDC42 inhibitors blocked vessel branching in the skin and colon of mice without apparent tissue toxicity. Taken together, these studies identify these agents as novel therapies for diseases characterized by disordered angiogenesis in skin or colon.

## Results

### RhoJ deletion disrupts the vascular network in the skin of RhoJ knockout (KO) mice

RhoJ KO mice have decreased branching of the retinal vasculature during development, a phenotype that improves with age (*26*). RhoJ is also known to play a role in adult vascular homeostasis, as RhoJ KO mice had impaired tumor angiogenesis (*36*) and impaired skin wound healing (*41*). However, the effects of RhoJ deletion on vascular homeostasis in adult skin has never been measured. Initial work sought to examine if RhoJ KO mice had altered vascular architecture in the skin as compared to RhoJ wild-type mice. We previously combined tissue clearing with intravascular infusion of DyLight 649 labeled tomato-lectin to visualize normal vasculature in the brain (*42, 43*). Here we modified an existing iDISCO (**i**mmunolabeling three- **D**imensional **I**maging of **S**olvent-**C**leared **O**rgans) (*44*) protocol to clear skin tissue harvested from RhoJ KO and wild-type mice infused with Tomato-lectin prior to sacrifice (Fig 1A). Cleared skin was imaged with a Leica SP8 multiphoton microscope to generate Z-stack images of skin and measure differences in vessel architecture between wild-type and RhoJ KO mice using two different analytic pipelines. A gross comparison of flattened images generated from RhoJ KO and wild-type mouse skin revealed that RhoJ KO mice had a decrease in the number of skin vessels (Fig 1B). Differences in vascular architecture in the skin were quantified by two different methods. AngioTool is a software package that takes 2D images, quantifies pixels in grayscale, and measures junctions and endpoints. This package has been used to measure differences in the vascular architecture of the murine embryonic hindbrain, post-natal retina, and allantois explant (*45*). NeuTube, in contrast, takes 3D z-stacks to trace neurons and can quantify terminal nodes, branch nodes, and the total number of neurons (*46*), and has only been recently applied to trace vessels. We compared the ability of these two methods to detect differences in vessel architecture in RhoJ KO (RhoJ-KO) and wild-type (RhoJ-WT) skin. AngioTool analysis revealed that the skin of RhoJ-KO mice had fewer vessel junctions and endpoints as compared to RhoJ- WT skin (Fig 1C). Analysis of the same images with neuTube revealed that RhoJ-KO skin had fewer terminal nodes, branch nodes, and vessel branches as compared to RhoJ-WT skin (Fig S4A-C). Despite the observed decrease in vessel density, RhoJ-KO skin showed no obvious pathological changes in the epidermis and dermis as visualized by haematoxylin and eosin staining (H&E) (Fig 1D, *Right*). We did, however, observe a slight decrease in thickness of the superficial dermal layer of RhoJ-KO skin, as highlighted by a Verhoeff-Van Gieson (VVG) stain (Fig 1D, *Left*). Flow cytometry analysis revealed that RhoJ-KO skin and RhoJ-WT skin had similar number of endothelial cells (CD31+ cells (*47*)) and fibroblasts (PDGRα+ cells (*48*)) in their skin (Fig 1E, *Top*) and spleen (Fig 1F, *Bottom*). Taken together, these results indicate that while RhoJ deletion affects vascular arborization in the skin, it does not grossly affect skin structure or the viability of fibroblasts or endothelial cells. Of note, three-dimensional quantitative approaches were able to better capture differences in the arborization of the vasculature in mice, and this approach was used to measure changes in vessel arborization in tumors, which are thicker structures than the skin. These studies suggest that inhibiting RhoJ/CDC42 GTPases may be a method to inhibit vascularization in skin.

**Fig. 1:**
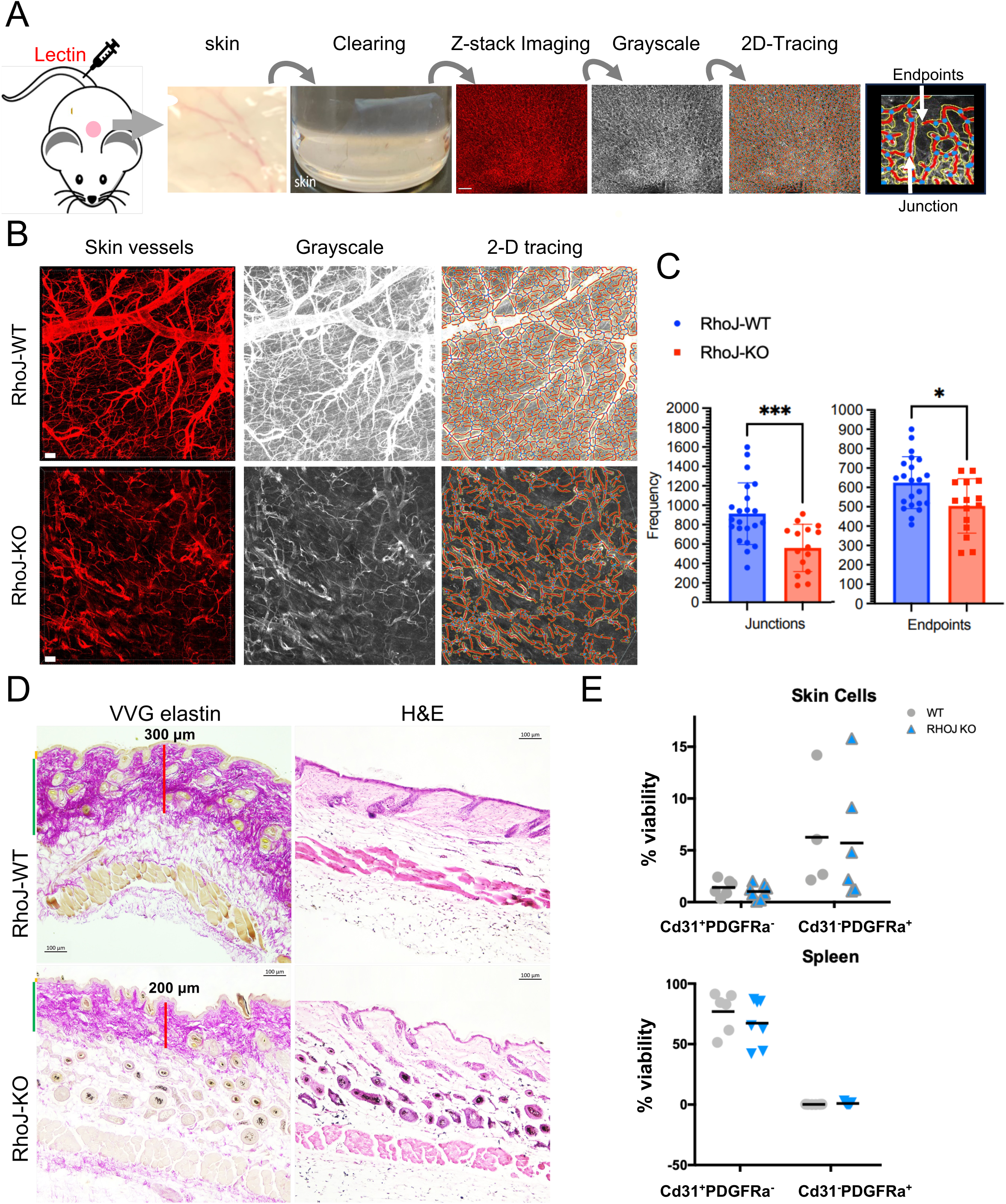
RhoJ regulates skin vessel arborization in skin. (A) Diagram of optical tissue clearing and vessel analysis. Each image (1.107 x 1.107 mm; 1024 x 1024 pixels) has about >50 z-stacks (5 μm/stack). Z-stacks are compressed into 2D Tiff file, converted to grayscale, and analyzed with AngioTool. Images were generated from three mice per group. (B) Representative images of skin vasculature from RhoJ wild type (WT) and knockout (KO) mice. Fluorescence images of lectin labeled structures were obtained, converted to grayscale, and traced with AngioTool. Scale bar is 50 μm. (C) Scatter bar plot of the frequency that vessel branches and termini were observed in (B); ^*,***^p < 0.013, 0.0009; n= 3 mice per genotype. (D) VVG elastin and H and E staining of skin from RhoJ-WT and RhoJ-KO mice. Note the presence of an intact epidermis and hair follicles indicating that RhoJ deletion did not induce skin necrosis. (E) Scatterplot showing the percentage of Cd31^+^ or Pdgfra^+^ live cells in the skin and spleen from >4 mice per group (RhoJ-WT and RhoJ-KO).

### CDC42 inhibitors alter vascular arborization patterns in patient-derived xenografts implanted into mice

Previous work discovered a new class of CDC42 interaction inhibitors that could inhibit tumor growth *in vivo* (*39, 40*) and block angiogenesis in vascularized microtumors *in vitro* (*40*). Initial work sought to examine whether the lead compound in our newly discovered class of CDC42 inhibitors (ARN22089) (*40*) could inhibit the growth and vascularization of melanoma patient-derived xenografts. A melanoma patient-derived xenograft was implanted into the flanks of NSG mice and when the tumors reached a volume of 150-200 mm^3^ they were treated with the indicated doses of the CDC42 inhibitor ARN22089, vemurafenib, or vehicle twice daily for two weeks by oral gavage (Fig 2A). We observed that ARN22089 inhibited the growth of tumors as effectively as vemurafenib. To compare how ARN22089 and vemurafenib affected arborization of vessels in tumors, we: 1) sacrificed the mice after the two week treatment; 2) infused the mice with tomato lectin intravascularly; 3) cleared the tumors using our iDISCO protocol; 4) imaged these tumors with multiphoton microscopy to obtain three-dimensional image stacks; 5) converted 3D images to grayscale images; and, 6) quantified vascular branching using neuTube (Fig 2B). Gross examination of stacked images from drug-treated mice revealed a decrease in vasculature in treated mice as compared to controls (Fig 2C, *Top*). Upon neuTube tracing of the vessel architecture (Fig 2C, *Below*), ARN22089 affected vessel number and branching to a greater extent than vemurafenib. NeuTube analysis of images revealed that ARN22089 had a dose-responsive effect on the number of branches, the number of terminal nodes, vessel tortuosity, and vessel length (Fig 2D). Analysis of the pathology of treated tumors revealed that while vemurafenib induced the accumulation of necrotic tissue, highlighted by hematoxylin, in treated tumor tissue (Figure 2E), ARN22089 treated tumors exhibited an expansion of the intracellular space (Figure 2E). We also performed daily tail vein injection of a second CDC42 lead inhibitor (ARN25062) and observed that this additional lead candidate: 1) stunted tumor growth (Fig S1A); and 2) reduced vessel arborization in tumors grossly (Fig S1B). Additionally, vessel arborization changes induced by these analogues could be detected using both AngioTool and neuTube (Fig S1C&D). Finally, we stained ARN22089 treated tumors with CD31, which revealed that treated tumors had fewer CD31-stained vessels (Fig 2F).

**Fig. 2:**
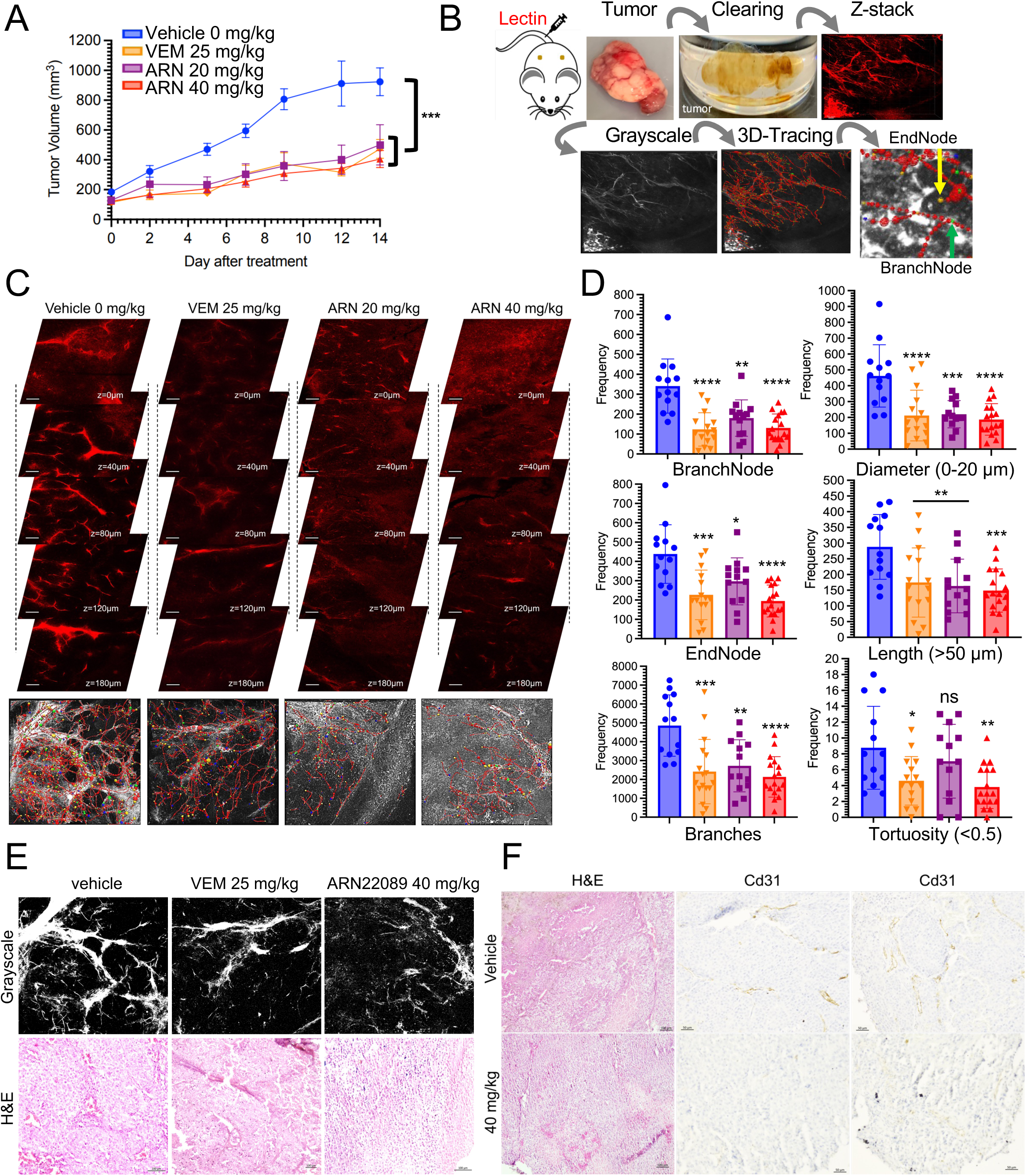
CDC42 inhibitors alter vascular arborization patterns in tumors. (A) Plot of tumor volume as a function of time in mice treated with vehicle (blue line), vemurafenib (orange line), or ARN22089 (purple [20 mg/kg] and red [40 mg/kg] lines) at the indicated doses. Graph represents mean+SEM. *p-value < 0.03 **p-value < 0.0071; ***p-value < 0.0001 (n=5 mice per group). (B) Schematic diagram of tissue clearing, visualization, and vessel analysis of the tumors. Each image (1.107 x 1.107 mm; 1024 x 1024 pixels) has >200 z-stacks (5 μm/stack). (C) Representative z-stacks of fluorescent images from tumors in mice treated with vehicle, vemurafenib (25 mg/kg twice daily) or ARN22089 (20 and 40 mg/kg twice daily) orally are shown. Bottom represents corresponding grayscale vessel tracing images; scale is 100 μm. (D) Scatter plot with bar graphs show the statistical quantification of vessel parameters (number of branch/end points and branching, average diameter, length, and tortuosity) after 3D vessel tracing for >3 tumors per group. ^*,**,***,****^p < 0.02, 0.003, 0.0006, 0.0001. (E) Representative H and E of vehicle, vemurafenib (25 mg/kg) and ARN22089 (40 mg/kg) tumor treated. (F) Representative H&E and CD31 staining of tumors showing a decrease in vessel-stained structures in tumors that were ARN22089 treated.

### CDC42 inhibitors block angiogenesis in human-derived vascularized microtumors *in vitro*

Previous work had demonstrated that ARN22089 could inhibit vessel growth in human derived vascularized microtumors *in vitro* using single-chamber **V**ascularized **M**icro**T**umor (VMT) models (*49*). Tumor cells in these models maintain: 1) their *in vivo* gene expression profile; 2) physiologic cell-cell interactions; and, 3) responsiveness to anti-cancer drugs (*50, 51*). To determine whether CDC42 inhibitors had direct effects on tumor vasculature, we used a second VMT platform design with independently treated vascular chambers and an intervening tumor chamber (Fig 3A) so that one could measure the effects of drugs after infusion on one side of the chamber versus the infusion of the other with vehicle. ARN22089 disrupted the vasculature in drug-treated but not vehicle treated chambers with a concomitant effect on tumor growth (Fig 3B), similar to the effects that were observed previously (*40*). We noted that ARN22089-treated chambers had shorter vessels (Fig 3C) as measured by REAVER (MATLAB)(*52*). Next, we sought to examine whether ARN25062 and vemurafenib could also inhibit vessel growth in a dual-chamber microfluidic device that consists of a vascular network adjacent to a second chamber containing the tumor, modeling ingrowth/cooption of vessels by tumors (Fig 3D). In the dual-chamber VMT, A375 tumors and associated vascular structures regressed significantly in response to treatment with ARN25062 and vemurafenib (Fig 3E), as indicated by a reduction in tumor growth on days 8 and 10 and decreased vessel length at each time point (Fig 3F). Notably, there was no significant difference in response between ARN25062 and vemurafenib in the VMT. Similar results were observed for the melanoma cell line WM3248, with a significant reduction in tumor growth and vessel length with ARN25062 and vemurafenib treatment, but only a significant decrease in vessel length with ARN25062 treatment (Fig 3G). Taken together, these results confirm that CDC42 inhibitors block the arborization of either mouse or human derived tumor vessels.

**Figure 3.**
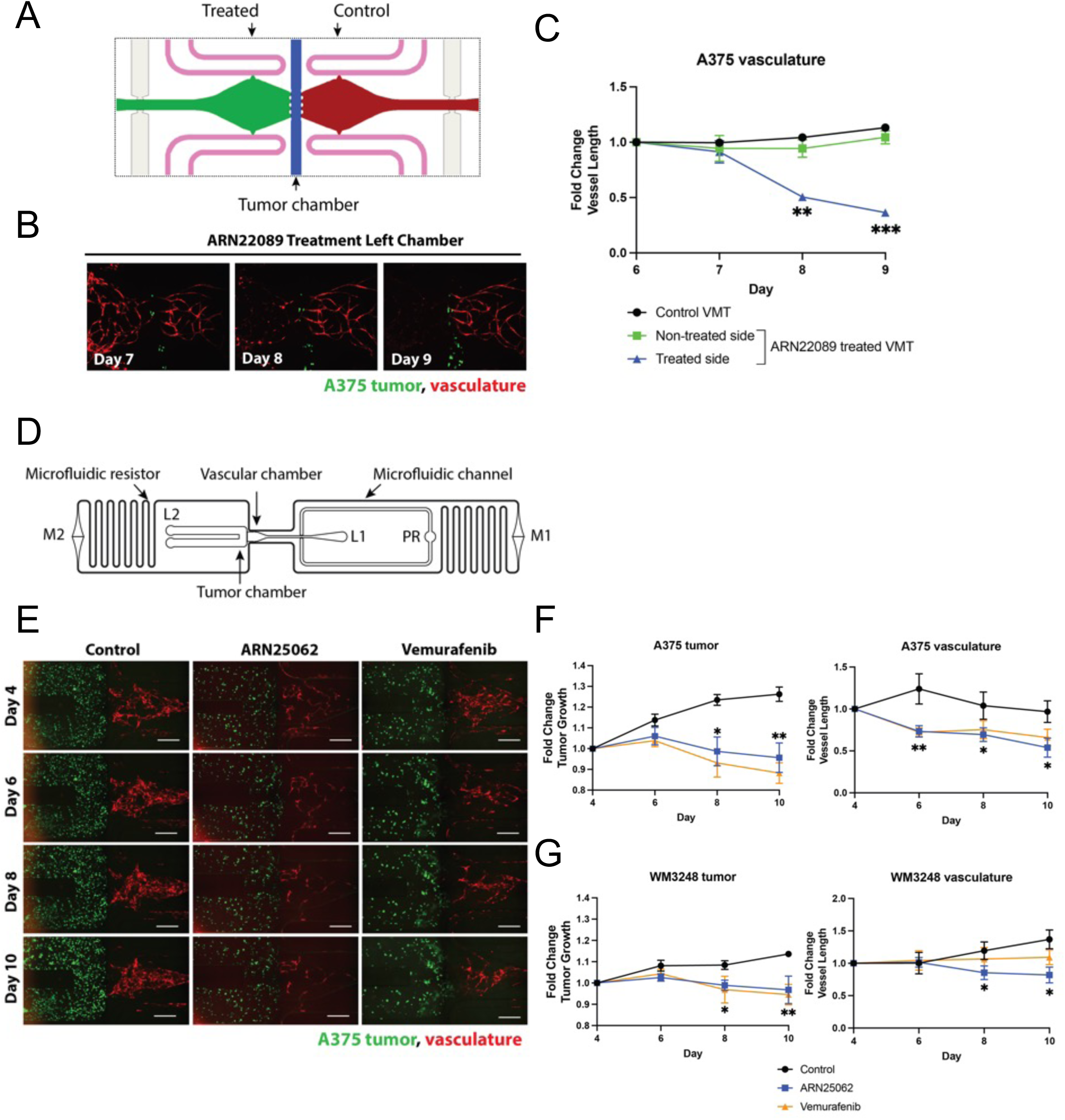
CDC42 inhibitors disrupt vasculature in tumor organoids. A) Diagram showing a double chamber microfluidic design where two independently fed vascular chambers (green and red) flank a central 200 µm wide tumor chamber (blue) separated by 3 PDMS posts on each side. The left vascular chamber (green) was treated with ARN22089 (2 µM) on day 6, while the right side received vehicle only (control, red). B) Representative fluorescent micrographs of double chamber VMT with A375 (green) in the center and vasculature (red) on either side. C) Quantification of A375 tumor-associated vasculature showing fold change in vessel length from baseline for left side (treated) vs. right side (control). D) Schematic showing a single dual-chamber microfluidic device. The vascular chamber connects to the tumor chamber (800 µm wide), separated by 6 PDMS posts spaced 50 µm apart that serve as burst valves to prevent the gel from traversing the chamber. EC and LF are introduced into loading port L1, and cancer cells are introduced separately into loading port L2. Loading is facilitated by a pressure regulator (PR). Tissues are maintained via hydrostatic pressure generated across microfluidic channels connecting media reservoirs M1-M2. Physiological flow rates are established by microfluidic resistors. E) Representative fluorescent micrographs of dual-chamber VMT containing A375 tumor (green) and vasculature (red), treated with either control (vehicle only), ARN25062 (2 µM), or vemurafenib (2 µM) for 48 hours starting on day 4. Media was refreshed on day 6, day 8, and day 10. Scale bar = 500 µm. F) Quantification of A375 tumor growth and tumor associated vasculature in the VMT showing fold change from baseline *p < 0.05, **p < 0.01. G) Quantification of WM3248 tumor growth and tumor-associated vasculature showing fold change in tumor vessel length from baseline. *p < 0.05, **p < 0.01.

### CDC42 inhibitors disrupt skin vasculature in mice in vivo and in human-derived vascularized microorgans in vitro

RhoJ KO mice had notable changes in the vascular arborization of skin compared to wild-type mice (Fig 1), suggesting that inhibiting RhoJ/CDC42 inhibitors could affect skin vasculogenesis. We next compared the effects of ARN22089 or vemurafenib treatment on the arborization of vessels in the skin of wild type mice (Fig 4). For these experiments: 1) C57B6 mice were treated with either ARN22089 (4, 12, or 40 mg/kg), vehicle, or 25 mg/kg vemurafenib twice daily for one week; 2) mice were infused with Tomato-lectin intravascularly prior to sacrifice; 3) skin tissues were cleared and imaged with three-dimensional multiphoton microscopy in 3D; and, 4) changes in vessel arborization were quantified using AngioTool. Gross examination of image stacks revealed that CDC42 inhibitors altered skin vascular arborization patterns, which were even more obvious when examining flattened images (Fig 4A, S2B). AngioTool was used to quantify changes in branching in the 2D images, revealing that ARN22089 had a dose-dependent effect on the number of observed junctions and endpoints (Fig 4B). In contrast, vemurafenib had no effect on the number of observed junctions and endpoints (Fig 4B). ARN22089 did not have any gross effects on skin appearance (Fig S2A) but did have modest effects on the thickness of the superficial dermal layer as detected by VVG staining and standard H&E staining (Fig 4C). These effects were similar to what was observed in RhoJ KO mice.

**Fig. 4:**
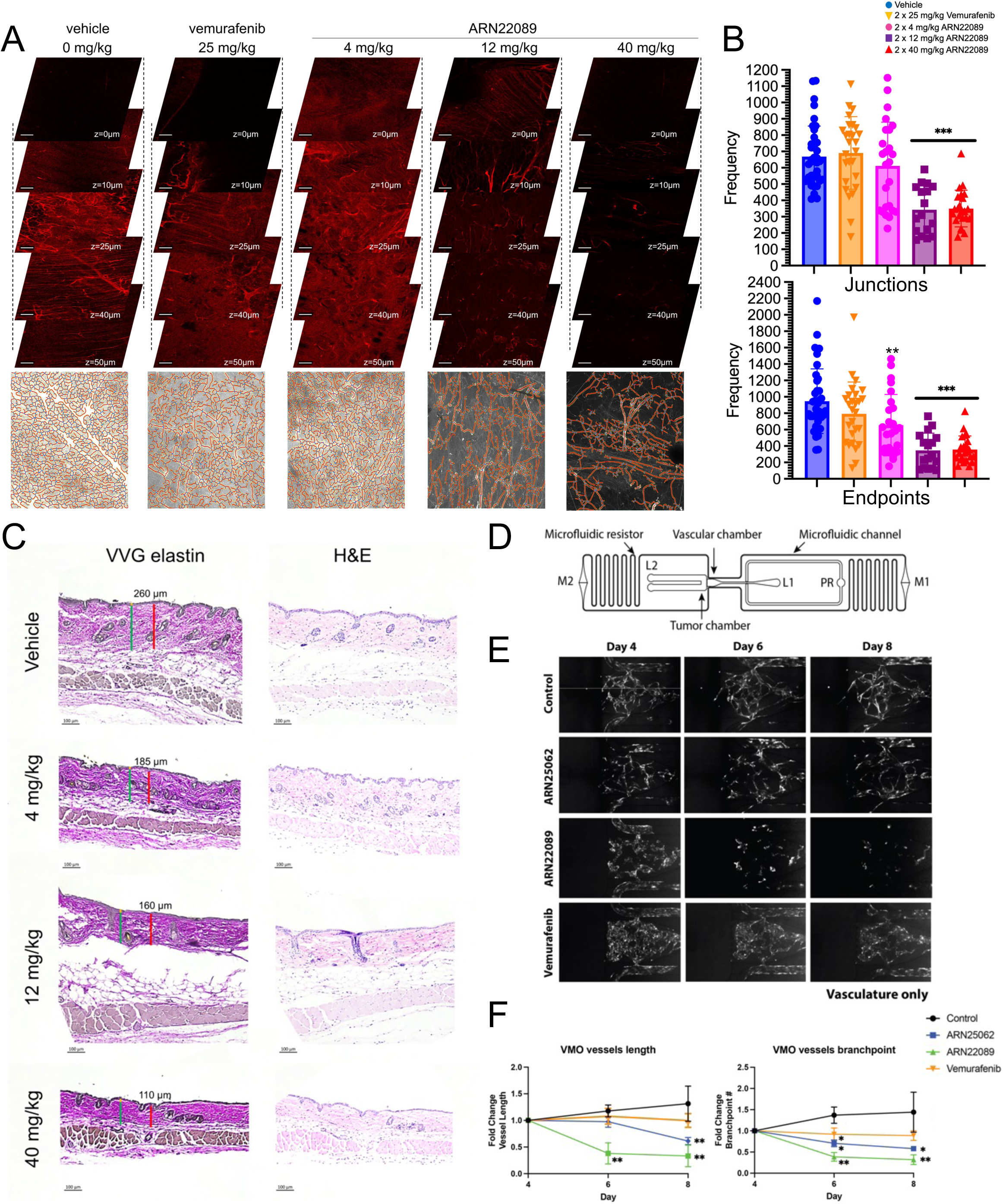
CDC42 inhibitors alter vessel arborization in skin. (A) Skin of mice treated with vehicle, vemurafenib, and ARN22089 at the indicated doses were cleared and imaged to generate Z-stacks as described, with representative stack shown. Bottom of each stack includes the 2D AngioTool tracing of pixels in grayscale. Scale bar is 100 μm. Each image (1.107 x 1.107 mm; 1024 x 1024 pixels) has >50 z-stacks (5 μm/stack). (B) Scatter plot with bar graphs show the frequency of branching and termini quantified from flattened images using the 2D AngioTool software. ^**,***^p-value < 0.0048, 0.0001, >3 mice per condition. (C) VVG elastin and H and E staining of skin from WT and RhoJ KO mice. Similar to the RhoJ KO, the presence of an intact epidermis and hair follicles are observed in all samples, with a slight decrease in VVG highlighted areas in treated animals. (D) Schematic showing a single dual-chamber microfluidic device. The vascular chamber is 800 μm wide, separated by 6 PDMS posts spaced 50 μm apart that serve as burst valves to prevent the gel from traversing the chamber. EC and LF are introduced into loading port L1. Loading is facilitated by a pressure regulator (PR). Tissues are maintained via hydrostatic pressure generated across microfluidic channels connecting media reservoirs M1-M2. Physiological flow rates are established by microfluidic resistors. (E) Representative fluorescent micrographs of dual chamber VMO showing vasculature (greyscale), treated with either control (vehicle only), ARN25062 (2 μM), ARN22089 (2 μM) or vemurafenib (2 μM) for 48 hours starting on day 4. Media was refreshed on day 6, day 8, and day 10. (F) Quantification of VMO-associated vasculature showing fold change in vessel length and number of branchpoints compared to baseline. *p < 0.05, **p < 0.01.

To further verify that ARN22089 affects skin vascularization, we harvested skin from the NSG mice bearing tumors that were treated with CDC42 inhibitors, vehicle, or vemurafenib at regions that were at a minimum 5 mm away from the implanted tumor. We observed that ARN22089 treatment at either 20 or 40 mg/kg BID inhibited the accumulation of vessel junctions and endpoints as measured by AngioTool (Fig S3A&B). Notably, vemurafenib did not significantly alter either the accumulation of vessel junctions or endpoints (Fig S3B), similar to what was observed in C57B6 skin.

Next, we examined how ARN22089, ARN25062, and vemurafenib affected vessels in **V**ascularized **M**icro-**O**rgans (VMO), which are the tumor-free version of the VMT micro- physiological system used above. These are comprised of human endothelial cells, pericytes, and fibroblasts, but no tumor cells (Fig 4D). We observed gross inhibition of vessel structure formation in the VMOs that were treated with ARN22089 or ARN25062, but this was not apparent in those treated with vemurafenib or vehicle (Fig 4E). REAVER quantification of VMO images revealed that treatment with ARN22089 and ARN25062 did indeed induce vascular disruption, evidenced by a significant reduction in vessel length and the number of branch-points (Fig 4F). In contrast, the vasculature was not significantly affected by vemurafenib treatment (Fig 4F).

To examine tissue toxicity, we next examined how ARN22089 affected the vascularization of other organs. ARN22089 had no effect on blood vessel accumulation in the brain (Fig S5A), although PK data did reveal that ARN22089 could cross the blood brain barrier to a small degree (Fig S5B). Haematoxylin and eosin staining of cardiac tissue harvested from treated animals revealed no gross effects on the heart (Fig S5C) and drug treatment had no effect on mouse weight (Fig S5D). Intriguingly, we did observe an effect on the accumulation of vasculature in the colon (Fig S5E) with changes in the appearance of villi (Fig S5E, *Below*).

### CDC42 inhibitors affect the vascularization of the skin by similar mechanisms observed in RhoJ KO mice

CDC42 inhibitors block the interaction of RhoJ and CDC42 with their downstream effectors (*40*), and both RhoJ KO mice and CDC42 inhibitor-treated mice had similar skin vessel arborization patterns. To better understand how ARN22089 modulates angiogenesis, we performed bulk RNA sequencing on WT skin treated with 12 mg/kg ARN22089 or vehicle twice daily for a week. We identified genes involved in CDC42 signaling and cell adhesion, as well as genes that mark cells carried within vascular compartments as genes that were significantly downregulated after drug treatment (Fig 5A). Notable amongst these genes were several with known roles in angiogenesis, including CDC42(*53*), RhoA(*54*), and CCL4 (*55*). Several genes involved in tumor angiogenesis/vasculogenesis, including leupaxin (*56*), CD177(*57*), and CCL12 (*58*), were also identified. Markers of lymphatic vasculature (Lyve1)(*59*) and genes that mark cell types trafficked in lymphatics (lymphocytes, neutrophils, macrophages) were also observed to be downregulated in drug treated skin (Figure 5B). Several components of PI3 kinase signaling were downregulated, consistent with earlier results that drug treatment affects S6 signaling (*40*). Importantly, we did not observe any gross skin toxicity in drug treated animals, as the structure of the hair follicles was preserved (Fig 4C) and there were similar numbers of fibroblasts and endothelial cells in the skin of drug treated and vehicle treated mice (Fig S2C). There was a slight decrease in the thickness of the VVG stained layer in the superficial dermis in drug treated mice (Fig 4C) as compared to control mice (Fig 1D), similar to what was observed in RhoJ KO mice.

**Fig. 5:**
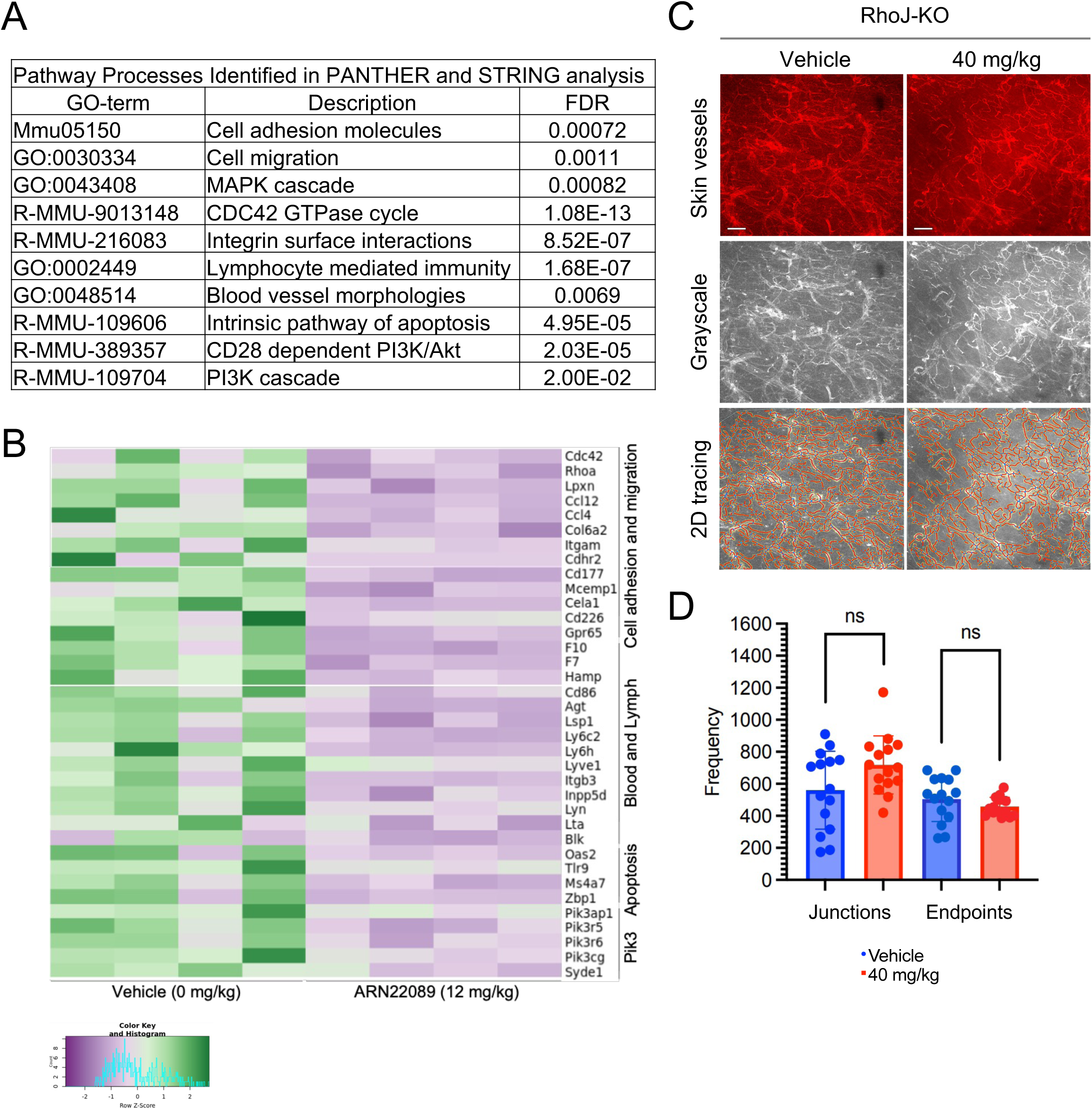
CDC42 inhibitors block skin angiogenesis in a RhoJ dependent manner. (A) Four mice were treated with 12 mg/kg ARN22089 or vehicle twice daily for one week, skin was harvested, and RNA was extracted from skin and subjected to bulk RNA sequencing (n=4 animals per group). Differentially expressed genes (p<.009) were used to identify pathways in the STRING and PANTHER database that were downregulated in ARN22089 treated skin (pathways with a False Discovery Rate < 0.002 are shown). (B) Heatmap showing significantly downregulated genes (at least 1.5-fold difference and p<.005) from the pathways identified in A. (C) Representative grayscale and AngioTool tracing of RhoJ KO skin vessels (treated with vehicle or 40 mg/kg ARN22089). (D) Scatter plot showing the frequency of branches and termini as measured by the 2D AngioTool software; n.s. no significant difference (n≥3 per condition).

As noted previously, both CDC42 and RhoJ have overlapping functions in skin angiogenesis and are required to integrate attractive and repulsive cues required for angiogenesis downstream of Plexin and VEGF receptors (*28*). To verify that ARN22089 blocked angiogenesis by inhibiting RhoJ/CDC42, we treated wild-type and RhoJ KO mice with ARN22089 and used the same approach as described in Figure 4 to measure changes in vascularization. Analysis of stacked and flattened images from the treated mice revealed no apparent differences in the vessel arborization patterns of RhoJ KO mice that were treated with ARN22089 from baseline (Fig 5C). Similarly, we observed no differences in junctions or endpoints between RhoJ KO treated and untreated mice as measured by AngioTool (Fig 5D). 3D vessel analysis with neuTube similarly observed no differences in the number of branch nodes and end nodes between RhoJ KO mice and RhoJ KO mice treated with ARN22089 (Fig S4A-C). NeuTube analysis also determined that there were more branch nodes and an increase in vessel length of wild type mice as compared to either KO mice or wild type mice treated with 12 mg/kg ARN22089 (Fig S4C). Taken together, these results are consistent with a model in which both CDC42 and RhoJ are critical regulators of blood vessel progression and regression in skin and tumors, such that inhibiting one or both of these pathways can alter the dynamics of blood vessels and affect vessel arborization in skin and tumors.

## Discussion

Existing methods to measure changes in vasculature rely on 2D quantitative approaches (*60, 61*), although several different 3D approaches have recently been used that integrate tissue clearing to measure tumor vascularization (*14, 16, 17*). We observe that a 3D quantitative approach (neuTube) provides more insight into vascular structure than approaches where 3D images are flattened into 2D. We show that this simple and straightforward approach used to trace neurons can be applied to measure changes in vessel arborization in tumors.

While many anti-angiogenic agents have been identified to treat cancer and other conditions, all of the existing agents have significant and frequent side effects in nearly half of treated patients, such as skin rash, diarrhea, and hypertension (*62, 63*). The cumulative side effects of these therapies preclude their use in conditions that arise in infancy, such as hereditary hemorrhagic telangiectasia, or in chronic diseases that require continuous treatment, such as inflammatory skin diseases. Moreover, the effects of VEGF inhibitors on vasculature are transient, as the normal vessel structures can return even one or two days after treatment cessation (*64, 65*).

Here, we sought to use our new imaging approach to examine how targeting angiogenesis pathways other than the conventional VEGF pathway, namely CDC42 GTPase-regulated pathways, affects angiogenesis. In contrast to the transient effects of RhoJ deletion on retinal vascular angiogenesis (*34, 35*), we discovered that RhoJ KO mice had a persistent decrease in vessel number in adult skin, suggesting that RhoJ has an important role in skin vascular homeostasis (Fig 1C). RhoJ deletion did not affect the number of endothelial cells and fibroblasts in skin (Fig 1E), in contrast to the significant differences in vessel branching observed. This suggests that RhoJ deletion affects endothelial cell behavior rather than survival. It is important to note that while the observed effects can be seen in normal skin, deletion does not grossly affect skin structure as the epidermis was intact and hair follicles were observed in RhoJ-KO mouse skin.

CDC42 inhibitors downregulated the expression of genes involved in cell migration and adhesion, and inhibited the accumulation of genes that mark leukocytes. The effects of CDC42 inhibitors on the skin largely recapitulated the effects observed with RhoJ deletion. Moreover, the activity of these agents required the presence of a functional RhoJ as they did not inhibit angiogenesis in RhoJ-KO mouse skin (Fig 5C&D). CDC42 inhibitors did not affect the vasculature of the heart or the brain but did affect vasculature in the colon (Fig S5A&C). It is known that vessel regression, which is regulated by RhoJ (*29, 66*), is important in the maintenance of the vascular plexus in the skin (*5*). Whether the observed tissue-specific differences are a consequence of different roles/expression patterns of RhoJ in distinct endothelial populations will be a topic for further study. Of note, other studies have shown that inhibiting receptors that activate RhoJ (Glutamate synthetase) can block inflammation in the skin induced by imiquimod (*41*), which further supports the role of RhoJ in skin vascular homeostasis. We observed that the effects in mice were preserved in human vascular organoids (Fig 4D-F), indicating that human blood vessels were also responsive to these drugs. The observation that CDC42 inhibitors can block angiogenesis in both mice and vascular organoid models suggested that CDC42 inhibitors could be used to treat conditions characterized by increased skin vascularity, such as rosacea, psoriasis, and photoaging. These agents may also be appropriate in treating conditions characterized by pathologic angiogenesis (port-wine birthmarks, hereditary hemorrhagic telangiectasias).

Melanoma tumors are minimally responsive to anti-angiogenesis agents (*6*). Tumor vessels can rapidly regrow after the cessation of VEGF-targeted therapies (*64, 65, 67*), and VEGF inhibitors seem to affect the blood vessels in the periphery but not the central part of the tumor (*17*). Here we used a combination of antibody administration, tissue clearing, and multiphoton microscopy to compare the effects of established melanoma therapies (BRAF inhibitors) and newly discovered CDC42 inhibitors in tumors. CDC42 inhibitors normalized vessel tortuosity and decreased vessel number, as did vemurafenib, a finding that has been reported for VEGF inhibitors (*17*). A closer examination of the histology of tumors revealed that vemurafenib-treated tumors were characterized by large areas of tissue necrosis, in contrast to CDC42 inhibitors where such broad necrosis was not observed (Fig 2E and Fig S1B, *Bottom H&E*). Vessel tracing results suggest that CDC42 inhibitors also affect large vessels in the tumor, in contrast to vemurafenib where large vessels were mostly preserved (Fig 2C). Moreover, vemurafenib did not affect skin vasculature, in contrast to CDC42 inhibitors. These observations suggest that vemurafenib likely affects vessel number by inducing the death and subsequent necrosis of melanoma cells with secondary effects on vasculogenesis, in contrast to CDC42 inhibitors which inhibit vasculogenesis directly in both skin and tumors.

In summary, the work presented here provides evidence that CDC42 GTPases play differing roles in vascular homeostasis in adult tissues. It is currently not clear whether the effects of CDC42 inhibition will be more permanent than VEGF inhibitors, yet it is known that bevacizumab does cross the blood brain-barrier and could induce some vascular toxicity in the brain. The relative lack of systemic toxicity in the brain and heart of CDC42 inhibitors compared with the rather specific effects of these agents on the skin and colon suggest that these agents could be used, at a minimum, as topical therapies for skin cancer and skin vascular/inflammatory disease.

## Materials and Methods

### In vivo skin and tumor experiments

All animal experiments were approved by the UC Irvine Institutional Animal Care and Use Committee (IACUC) (AUP-20-161). C57BL6 mice and RhoJ KO mice in the C57B6 background were used in studies examining skin, colon, and brain vascularization while NOD.Cg-*Prkdcscid IL2rgtm1Wj*/SzJ (NSG) mice were used for the patient-derived xenograft experiments. An equal number of male and female mice were used in treatment with inhibitors. For skin treatment, wild-type or RhoJ KO C57BL6 mice were treated with inhibitors by oral gavage twice daily for one week. For PDX tumor experiments, tumors were maintained by passaging into NSG mice before treatment. Inoculation of tumors in NSG mice was performed as previously described (*49*). When tumors reached ∼150-200 mm^3^, mice were treated by oral gavage twice daily with 20 or 40 mg/kg ARN22089 or with vemurafenib at 10 or 25 mg/kg. Tail-vein injection was done once a day at 10 mg/kg ARN22089. All tumor experiments involved treating animals for two weeks while skin experiments involved treatment for 1 week.

### Cardiac perfusion and tissue collection

At time of harvest, mice were infused with lectin-DyLight-649 (200 uL, 25% lectin-DyLight and 75% PSB) through the tail vein injection to label endothelial cells. One hour later, mice were euthanized by cardiac perfusion with 50 mL saline followed by 4% methanol-free paraformaldehyde (PFA) as described previously(*43*). For skin, the fur was shaved and depilated and dorsal skin was removed from the back. Tumors were removed from the flanks of the dorsal region. Brain, kidney, intestines, livers, hearts, and tissues were placed in 4% PFA for 24-48 h and then placed in 1xPBS at 4°C for subsequent tissue clearing.

### Tissue clearing and imaging analysis

A modified iDISCO protocol was used to clear tissues. Tissues were dehydrated in graded series (20, 40, 60, 80, 100, 100%) of methanol in water for 48 h at RT. Samples were incubated with 66% dichloromethane (DCM) and 33% methanol for 48 h at RT. Next, samples were washed twice for 15 min each with DCM and placed in dibenzyl ether (DBE) for storage and used as a medium for imaging. The Leica TCS SP8 X instrument was used to image tissues and processed with a LAS X Navigator suite. The sample was placed on a makeshift holder attached to a microscope slide and filled with DBE; a coverslip was placed over the sample, making sure the sample and liquid touches the coverslip. Images were captured using an HC PL FLOUTAR 10x/0.30 objective. Image dimension: 1024 x 1024 pixel dimension and 1107.14 μm by 1107.14 μm, 5 µm per z-stack. Vascular fluorescence was detected by scanning for signals in the DyLight649 spectrum (670-708 nm). For image analysis, Z-stack images were opened with Fiji ImageJ, and color images were converted to black background and white pixels using color-split and saved as a tiff file either in three-dimenasional (3D) or compressed into 2D image using Z stack projection. 3D tiff files were analyzed using neuTube software (*46*), while 2D images were analyzed using AngioTool software (*45*). Both automated and manual tracing were performed on Tiff images in neuTube. Automated tracing was applied initially, with manual curating as needed due to images with low contrast between vessels and background and where large vessels cannot be correctly auto-traced. A custom MATLAB script was used to extract the number of branch and end points, vessel length, vessel diameter, and tortuosity in the neuTube SWC format files (individual nodes with x, y, z coordinates, radius and node connectivity). In the SWC format, vessel structures are simplified into spherical nodes with the radius of the vessel at a given location. Branch points (green nodes in images) appear where branches occur, and end points (yellow nodes in images) appear at the end of a vessel segment (all other nodes are red in images). A vessel segment is defined as a series of nodes that are bounded by two branch points or one branch point and one end point. The length of a vessel segment is defined as the distance between its pair of boundary points. The diameter of the vessel is calculated using the radius of the nodes. Parameters for AngioTool analysis were set the same for all images analyzed. For NSG skin stack images, the first and last few images in the stacks were removed to reduce background noise; all stacks from C57BL6 skin were analyzed. For image analysis >3 tumors or skin tissues were cleared, imaged and analyzed.

### Statistics

GraphPad Prism software was used to generate graphs and perform statistical significance, using one- and two-way ANOVA test and unpaired T-test. One-way ANOVA and unpaired T-test were applied on vessel analysis and two-way ANOVA test was used to determine significance for tumor growth curves. Custom R script was used to convert pixel value to micrometer for diameter and distance; it was also used to calculate the frequency of branch nodes, end nodes, number of vessels and branching, and tortuosity.

### Microfluidic device fabrication

Microfluidic device fabrication and loading have been previously described (5–6). In summary, a custom polyurethane master mold was created using a two-part polyurethane liquid plastic (Smooth Cast 310, Smooth-On Inc.). Subsequently, a PDMS layer was replicated from this master mold, and holes were punched to create inlets and outlets. The platform was assembled in two stages: first, the PDMS layer was chemically glued and subjected to 2 minutes of oxygen plasma treatment to affix it to the bottom of a bottomless 96-well plate (Greiner). Following this, a 150 µm thin transparent membrane was bonded to the bottom of the PDMS device layer through an additional 2-minute treatment with oxygen plasma. The fully assembled platform was then placed in a 60°C oven overnight, covered with a standard 96-well plate polystyrene lid, and sterilized using UV light for 30 minutes before cell loading.

### Cell culture and microfluidic device loading

To establish the vascular chamber, normal human lung fibroblasts and ECFC-ECs (endothelial colony forming cell endothelial cells) or HUVECs (human umbilical vein endothelial cells) were harvested and resuspended in fibrinogen solution at a concentration of 3×10^6^ cells/mL and 7×10^6^ cells/mL, respectively. For VMT, A375 and WM3248 melanoma cells were introduced into the tumor chamber at a concentration of 1 × 10^5^ to 2 × 10^5^ cells/mL fibrinogen solution. Fibrinogen solution was prepared by dissolving 70% clottable bovine fibrinogen (Sigma- Aldrich) in EBM2 basal media (Lonza) to a final concentration of 5 mg/mL. The cell-matrix suspension was mixed with thrombin (50 U/mL, Sigma-Aldrich) at a concentration of 3 U/mL, quickly seeded into the microtissue chambers, and allowed to polymerize in a 37 ◦C incubator for 15 minutes. Laminin (1 mg/mL, LifeTechnologies) was then introduced into the microfluidic channels through medium inlets and incubated at 37 ◦C for an additional 15 minutes. After incubation, culture medium (EGM-2, Lonza) was introduced into the microfluidic channels and medium wells. The medium was changed every other day, and the hydrostatic pressure head re-established daily to maintain interstitial flow.

### Drug treatment in the VMO and VMT

Following a culture period of 4–5 days to facilitate the development of a perfused vasculature within each VMO or VMT, the culture medium was replaced with a medium containing the specified drug concentrations. Drugs were administered to the microtissues through the newly formed vascular bed via gravity-driven flow. Specifically, ARN25062, ARN22089, and vemurafenib were used at a 2 µM dose. A375 VMT, WM3248 VMT, and VMO were randomly assigned to one of four conditions: control (vehicle only), 2 µM ARN25062, 2 µM ARN22089, or 2 µM vemurafenib. For experiments with two vascular side chambers, the left chamber received 2 µM ARN22089, while the right chamber served as control (vehicle only). Both VMO and VMT underwent a 48-hour treatment period, with complete medium replacement every 48 hours. Fluorescent micrographs of VMT were captured every 48 hours for 6 days post-treatment, and the quantification of tumor and vasculature growth was performed.

### Fluorescence imaging and analyses

Fluorescence images were acquired with a Biotek Li- onheart fluorescent inverted microscope using automated acquisition and standard 10x air objective. To test vessel perfusion, 25 µg/mL FITC- or rhodamine- conjugated 70 kDa dextran was added to the medium inlet prior to treatment. For quantifying vessel length in VMOs and VMTs, REAVER software (MATLAB (7)) was employed. ImageJ software (National Institutes of Health) was utilized to determine the total fluorescence intensity (mean grey value) for each tumor image, providing a measure of tumor growth. Normalization to baseline was performed for each chamber. In VMTs, tumor growth was quantified by measuring the total fluorescence intensity in the color channel representing the tumor cells. This measurement accounted for both the area and depth of individual tumors, considering that thicker areas appear brighter. Any image adjustments made were applied uniformly to ensure consistency across all images in the experimental group.

### Tissue Histology and Immunohistochemistry

Tissues were fixed in 4% formaldehyde and washed in 1xPBS. For embedding, tissues were incubated in 65% ethanol for 30 min at RT and kept in 70% ethanol before sending the samples to Experimental Tissue Resource (ETR) at UCI for embedding, VVG (*68*) and H&E staining (*69*).

### Flow Cytometry

Back of mouse was first shaved and depilated. A large surface area (6 cm x 3 cm) of the skin on the back was cut and fat removed. The skin was minced, and digest buffer added (RPMI without Ca/Mg (no FBS/EDTA), 23.2 mM HEPES and 2.32 mM sodium pyruvate, 0.25 mg/mL Liberase, 46.4 unit DNase) and incubated for 1.5 h at 37°C shaking. Spleen samples were removed from the animals, minced and RPMI added to minced tissue. All samples were filtered with 70 micrometer cell strainer, and washed with FAC buffer (5% FBS, 2 mM EDTA, 1% non-essential amino acids, 3.9x beta-mercapthoethanol (from 1000x)) twice and spun at 18.0 xg for 10 min 4°C. Skin samples were blocked with TruStain FcX PLUS and spleen samples with Fc block, both, at 1:100 for 15 min at 4°C. All samples washed twice with FAC buffer and incubated in primary Cd31 (1:50, brilliant violet 421 Invitrogen 404031182), PDGFRa (1:100, APA5 Life Technology 12140181), viability (1:200, eBioscience dyefluor 780 65086514) in FAC buffer for 30 min at 4°C nutating. Samples were then washed 2x and fixed with 1% formaldehyde for 15 min and washed twice again. Samples were resuspended with 400 uL FAC buffer and ran on the BD Fortessa X20 Flow cytometer. Compensation and FMO for both skin and spleen were run before running the experimental samples. Samples were analyzed using FlowJo.

### Quantification of brain levels of ARN22089 following oral administration

Animals were treated with ARN22089 at 10 mg/Kg by oral administration, as described (39). Mouse brains were collected at 1, 2, 4 and 8 hours after administration. The brains were homogenized and, following protein precipitation with acetonitrile, the amount of ARN22089 was quantified by LC-MS/MS as described (39).

### Bulk RNA sequencing and pathway analysis

A small piece (approximately 3 x 3 cm section) of the skin from a mouse was removed and placed in Qiagen buffer RLT (plus beta-mercaptoethanol). A Precellys Homogenizer was used to homogenized the skin sample in Precellys beads. RNeasy kit (Qiagen) was used to extract RNA, with DNase I digestion step. The RNA samples were sent to the Genomic Core Facility at UCI for library construction and sequencing. Paired-end sequencing reads were aligned and feature count to the mouse reference UCSC/mm10 (ERCC spike-in reference was concatenated with the mouse genomic and transcript reference) (*70*)) with HISAT2 v2.2.1 (*71*) and samtools/1.15.1 (*72, 73*). Count matrix was generated in R programming and RUVSeq (*74*) package was used to normalize (RUVr, residuals) RNA-seq data and determine differential expression. List of differential genes (vehicle vs ARN22089 (12 mg/kg)) was used to run functional processes on PANTHER (*75, 76*) (p<0.09) and STRINGv12 (*77*) (p<0.009) to determine pathway analysis (*78*) and gene ontology (see Supplemental Tables). Heatmap was generated using R programming.

## Acknowledgement

Funding for this work was largely derived from (CA-244571) from the National Cancer Institute and S10OD028698. This study was made possible in part through access to the Optical Biology Core Facility, a shared resource supported by the Cancer Center Support Grant (CA-62203) and UCI Skin (AR-075047) at the University of California, Irvine.

## Author contributions

Conceptualization: AKG, CCWH, MDV, BC, LMV

Funding acquisition and project administration: AKG

Data curation: LMV, SJH, JS, SMB, NB, MS, RB, AA

Formal analysis: LMV, SJH, JS, SMB, NB, MS, RB, AA, VSHK

Investigation: LMV, SJH, JS, NS, DFX, TN

Writing – original draft: LMV, AKG

Writing – review and editing: LMV, AKG, CCWH, MDV, BC, DFX, SJH, JS

Methodology: NB, SMB, MS, RB, AA, TN, NS, DFX, RP, VSHK

