## Supplemental Figures for "CDC42 Inhibitors Alter Patterns of Vessel Arborization in Skin and Tumors in vivo"

### Supplemental Figure 1: ARN25062 affects tumor vascular architecture.

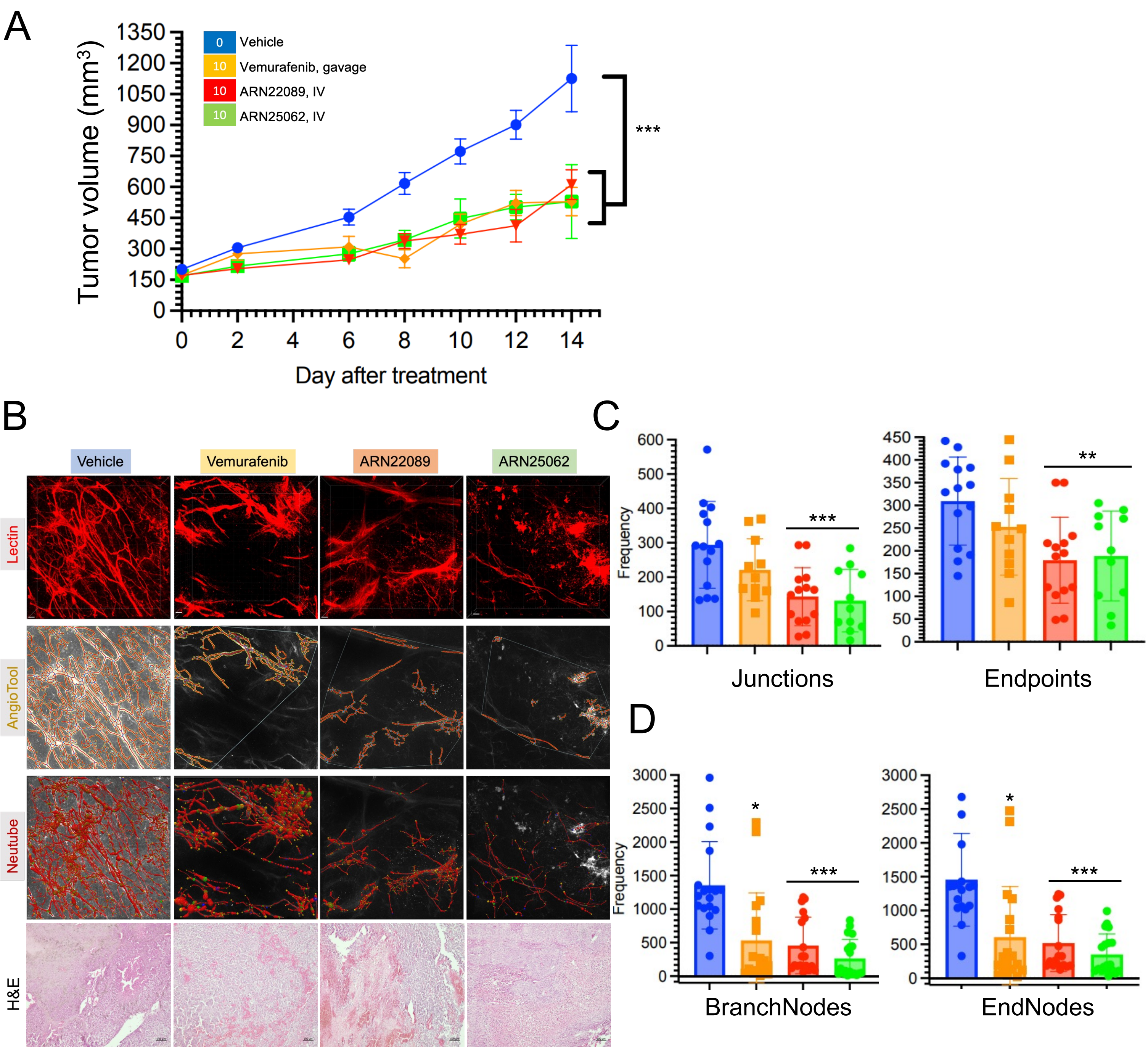

**Supplemental Figure 1. RhoJ deletion modulates skin vascular architecture.** (A) Diagram of optical tissue clearing and vessel analysis. Each image (1.107 x 1.107 mm; 1024 x 1024 pixels) has about 50-100 z-stacks (5 μm/stack). Z-stacks are compressed into 3D Tiff file, converted to grayscale, and analyzed with neutube. Images were generated from three mice per group. (B) Representative images of skin vasculature from RhoJ wild type (WT) [vehicle vs 12 mg/kg twice daily a week] and knockout (KO) [vehicle vs 40 mg/kg twice daily a week] mice with neuTube tracing. Fluorescence images of lectin labeled structures were obtained, converted to grayscale, and traced with neutube. Scale bar is 100 μm. (C) Scatter plot with bar graphs show the statistical quantification of vessel parameters (number of branch/end points and branching, and tortuosity) after 3D vessel tracing. Unpaired 2-tail T-test was used to determine p-value. $^{*,**}p < 0.03, 0.005$, at least 3 skin samples per condition were analyzed.

**Supplemental Figure 2: CDC42 inhibitors affect blood vessel arborization without inducing overt toxicity.**

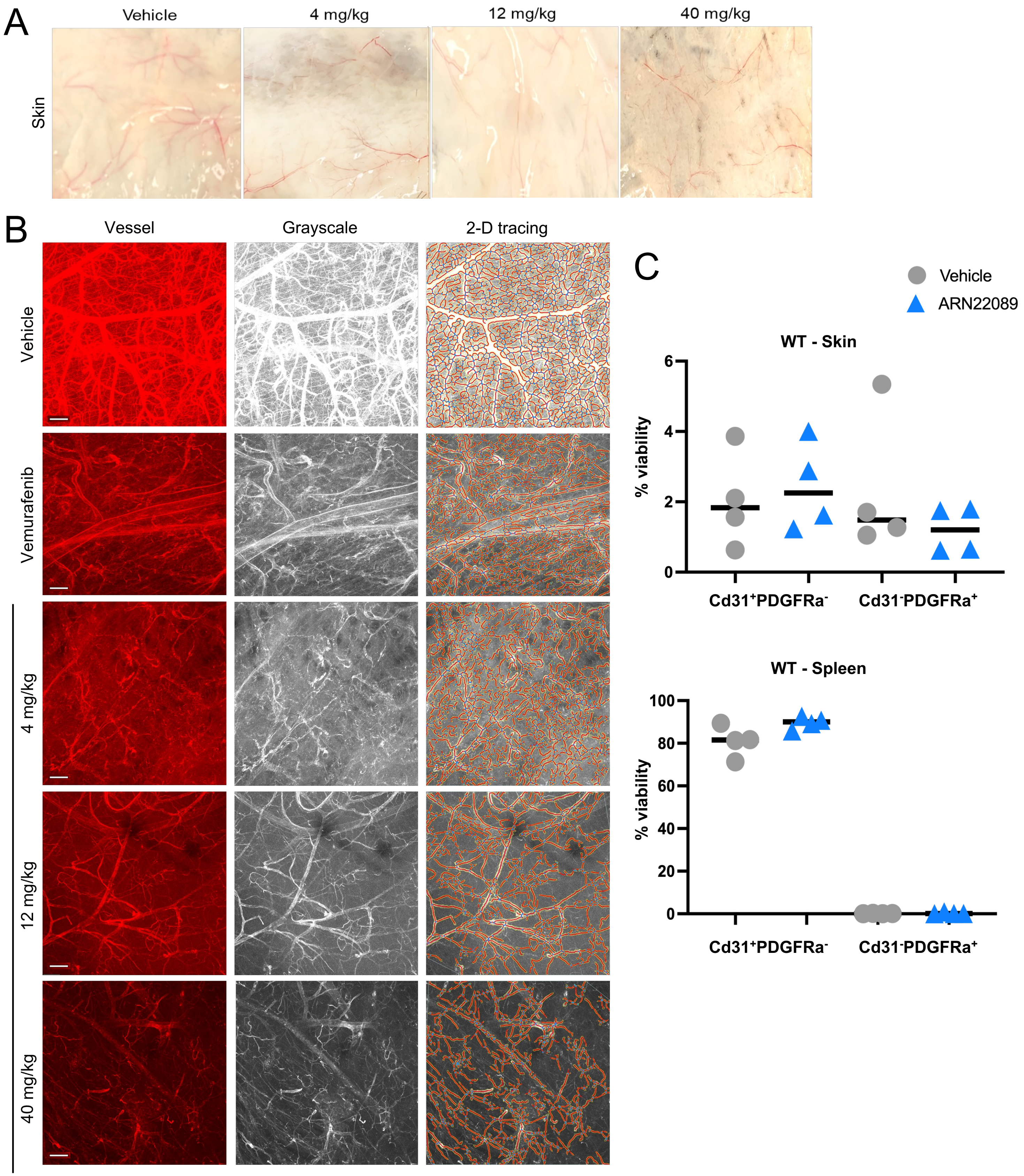

**Supplemental Figure 2: ARN25062 disrupts tumor angiogenesis.** (A) Growth curves of tumors treated with the indicated doses of ARN25062 and ARN22089 by IV as compared to administration via oral gavage of vemurafenib at 10 mg/kg BID once daily for two weeks. Mean±SEM, Two-way ANOVA test indicates significance between vehicle and treated mice for ARN22089, ARN25062 and Vemurafenib ***p < 0.0003. (B) Representative 3D images of cleared tumor showing vessels labeled with lectin dyLight from mice treated with indicated inhibitors. *Center*, images were converted to grayscale and saved as a tiff file for 2D tracing with AngioTool or 3D tracing with neuTube. *Bottom*, H&E staining of tumors. Scale bar = 50 µm. Each image (1.107 x 1.107 mm; 1024 x 1024 pixels) has about 207 z-stacks (5 µm/stack) (C) Scatter plot with bars show the statistical quantifications from Angiotool tracing of vessel branching and endpoints. (D) Scatter plot with bars show the statistical quantifications from Neutube tracing of vessel branching and endpoints. Unpaired 2-tail T-test applied in 2D and 3D analysis; $^{*,**,***}p < 0.002$, 0.006, 0.0002. At least 3 tumors per condition were analyzed.

**Supplemental Figure 3: ARN222089 inhibits angiogenesis in skin adjacent to tumors in NSG mice.**

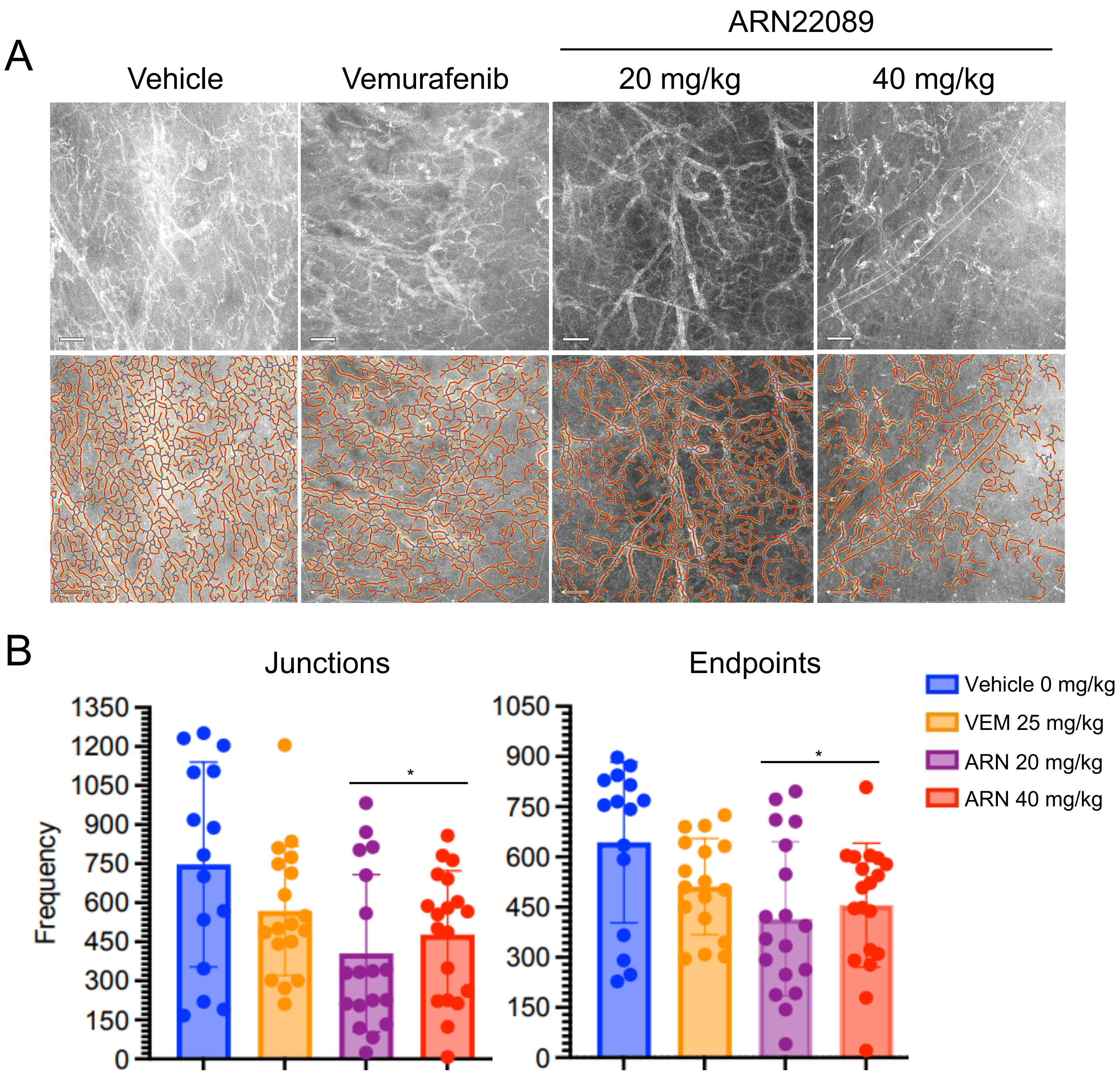

**Supplemental Figure 3: CDC42 inhibitors affect blood vessel arborization without inducing overt toxicity.** (A) Micrographs of skin harvested from treated mice. Note the presence of observable vessels. (B) Skin of mice treated with vehicle, vemurafenib, and ARN22089 at the indicated doses, optically cleared and imaged to generate Z-stacks. Flattened images from additional samples to those presented in Figure 4A are shown. Images were converted to grayscale and vessels traced with Angiotool. Scale bar is 100 μm. Each image (1.107 x 1.107 mm; 1024 x 1024 pixels) is from >50 flattened z-stacks (5 μm/stack). (C) Scatterplot showing percent live skin and spleen with $Cd31^{+}$ or $Pdgfra^{+}$ from 4 mice per group (RhoJ-WT: vehicle vs 12 mg/kg, n=4 per group)

**Supplemental Figure 4: RhoJ deletion modulates skin vascular architecture.**

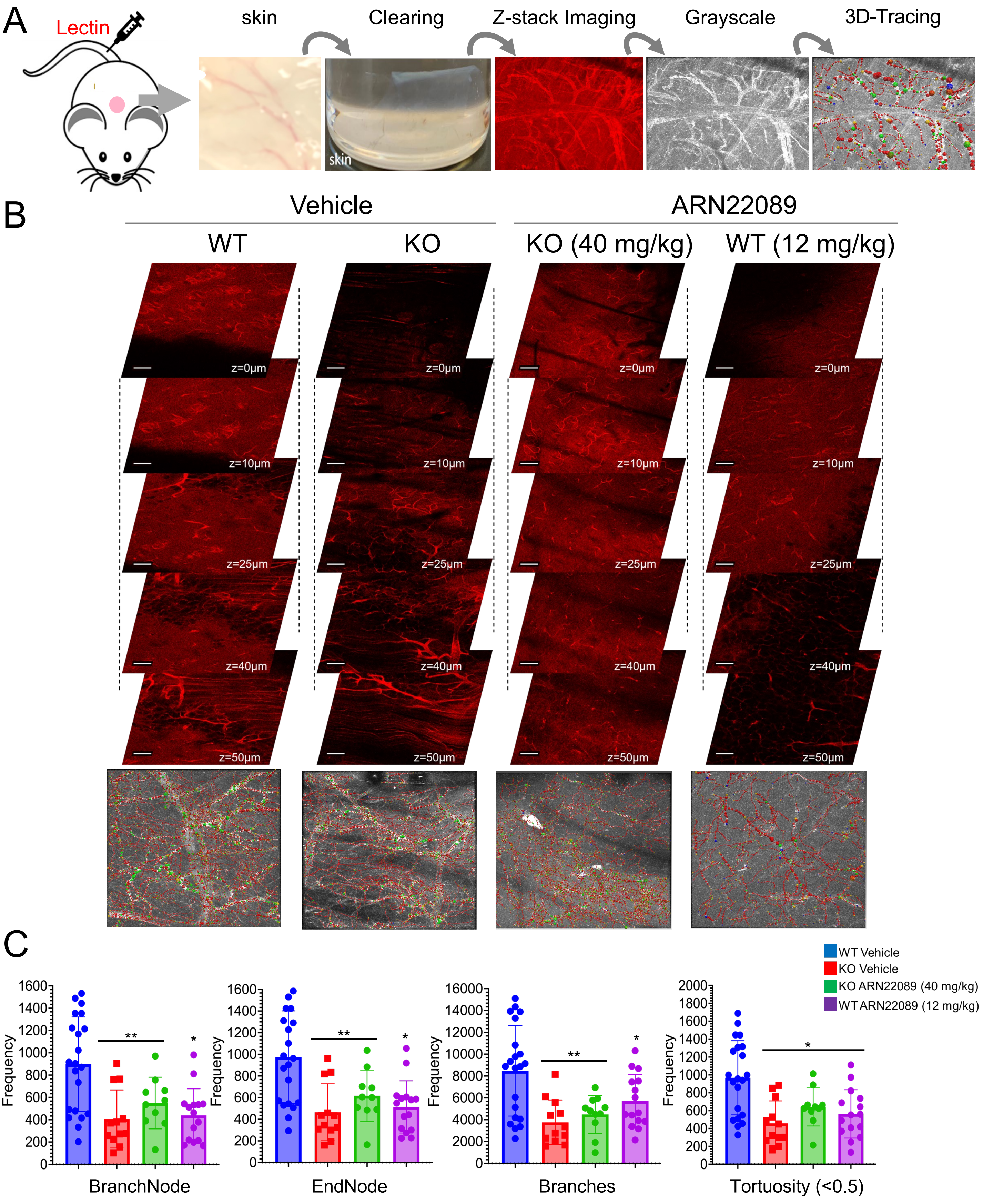

**Supplemental Figure 4: ARN222089 inhibits angiogenesis in skin adjacent to tumors in NSG mice.** (A) Representative grayscale images of adjacent skin vessels from mice bearing tumors that were treated with 20 or 40 mg/kg twice a day of inhibitor for two weeks. Scale bar = 100 µm. Below shows 2D tracing with Angiotool. (B) Scatter plots show number of branches and vessel termini in mice treated with 20 and 40 mg/kg of ARN22089. Each image (1.107 x 1.107 mm; 1024 x 1024 pixels) and has >50 z-stacks (5 µm/stack). $^{*}p < 0.02$ with at least 3 tumors per group.

**Supplemental Figure 5: ARN22089 disrupts vessel in skin and colon but not the brain.**

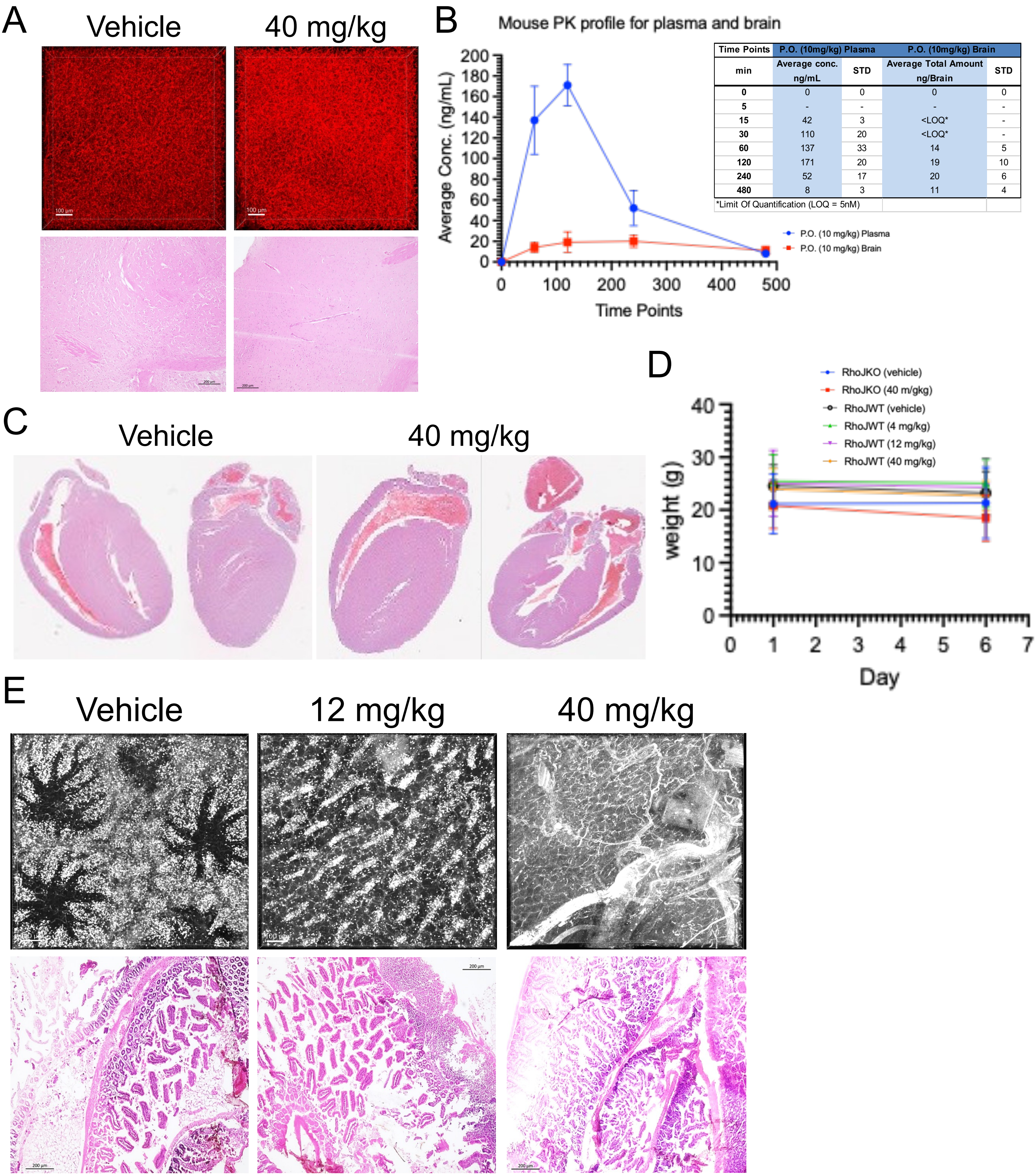


| Time Points | P.O. (10mg/kg) Plasma | | P.O. (10mg/kg) Brain | |
|---|---|---|---|---|
| min | Average conc. ng/mL | STD | Average Total Amount ng/Brain | STD |
| 0 | 0 | 0 | 0 | 0 |
| 5 | - | - | - | - |
| 15 | 42 | 3 | <LOQ* | - |
| 30 | 110 | 20 | <LOQ* | - |
| 60 | 137 | 33 | 14 | 5 |
| 120 | 171 | 20 | 19 | 10 |
| 240 | 52 | 17 | 20 | 6 |
| 480 | 8 | 3 | 11 | 4 |

*Limit Of Quantification (LOQ = 5nM)

**Supplemental Figure 5: ARN22089 disrupts vessels in skin and colon but not the brain.** (A) Representative images of brain vessels in red labeled with lectin dyLight. Scale bar = 100 μm, from mice treated with the indicated doses of ARN22089 or vehicle. Representative H&E staining of the brain tissue. (B) PK profile of ARN22089 in the plasma and brain after oral administration of 10 mg/kg. The amount of compound detected in the brain was about 10 fold less than that in the plasma. (C) H and E images of the hearts from mice that were treated with 0 mg/kg vehicle or 40 mg/kg ARN22089. (D) ARN22089 treatment does not affect mouse viability. Weights of mice treated with ARN22089 were measured after day and day 6 of treatment; mean$\pm$SD. (E) Representative images of cleared gastrointestinal tissues from mice treated with the indicated doses of ARN22089. Tissues were labeled with lectin dyLight, red fluorescence images were converted to grayscale using ImageJ. Decreased vessels were observed at the 12 and 40 mg/kg doses. Representative H&E staining of the intestinal tissues.

**Supplemental Table: Differential gene expression and pathway analysis from WT skin treated with ARN22089 at 12 mg/kg. Tab 1** includes the list of differential gene analysis from the bulk RNAseq pipeline and RUVseq package in R. **Tab 2** provides a complete name of the genes from Tab 1 ($p<0.09$) that were mapped to PANTHER pathway analysis. **Tab 3** shows the PANTHER overrepresentation test with reactome pathways from genes in Tab 2. **Tab 4** includes functional analysis identified in STRING (v12.0). Based on the pathways shown in Tab 3, the genes with $p<0.009$ were annotated in STRING to determine functional processes.
